# Benchmarking undedicated cloud computing providers for analysis of genomic datasets

**DOI:** 10.1101/007724

**Authors:** Seyhan Yazar, George E C Gooden, David A Mackey, Alex W Hewitt

## Abstract

A major bottleneck in biological discovery is now emerging at the computational level. Cloud computing offers a dynamic means whereby small and medium-sized laboratories can rapidly adjust their computational capacity. We benchmarked two established cloud computing services, Amazon Web Services Elastic MapReduce (EMR) on Amazon EC2 instances and Google Compute Engine (GCE), using publicly available genomic datasets (*E.coli* CC102 strain and a Han Chinese male genome) and a standard bioinformatic pipeline on a Hadoop-based platform. Wall-clock time for complete assembly differed by 52.9% (95%CI: 27.5-78.2) for *E.coli* and 53.5% (95%CI: 34.4-72.6) for human genome, with GCE being more efficient than EMR. The cost of running this experiment on EMR and GCE differed significantly, with the costs on EMR being 257.3% (95%CI: 211.5-303.1) and 173.9% (95%CI: 134.6-213.1) more expensive for *E.coli* and human assemblies respectively. Thus, GCE was found to outperform EMR both in terms of cost and wall-clock time. Our findings confirm that cloud computing is an efficient and potentially cost-effective alternative for analysis of large genomic datasets. In addition to releasing our cost-effectiveness comparison, we present available ready-to-use scripts for establishing Hadoop instances with Ganglia monitoring on EC2 or GCE.

## INTRODUCTION

Through the application of high-throughput sequencing, there has been a dramatic increase in the availability of large-scale genomic datasets.[1] With reducing sequencing costs, small and medium-sized laboratories can now easily amass many gigabytes of data. Given this dramatic increase in the volume of data generated, researchers are being forced to seek efficient and cost-effective measures for computational analysis.[2] Cloud computing offers a dynamic means whereby small and medium-sized laboratories can rapidly adjust their computational capacity, without concern about its physical structure or ongoing maintenance.[3–6] However, transitioning to a cloud environment presents with unique strategic decisions,[7] and although a number of general benchmarking results are available (http://serverbear.com/benchmarks/cloud; https://cloudharmony.com/), there has been a paucity of comparisons of cloud computing services specifically for genomic research.

We undertook a performance comparison on two established cloud computing services: Amazon Web Services EMR on Amazon EC2 instances and GCE. Paired end sequence reads of publicly available genomic datasets (*Escherichia coli* CC102 strain and a Han Chinese male genome) were analysed using Crossbow, a genetic annotation tool, on Hadoop-based platforms with equivalent system specifications.[8–10] A standard analytical pipeline was run simultaneously on both platforms multiple times (Figure 1 and Supporting Information Figure 1). The performance metrics of both platforms were recorded using Ganglia, an open-source high performance computing monitoring system.[11]

**Figure 1.**
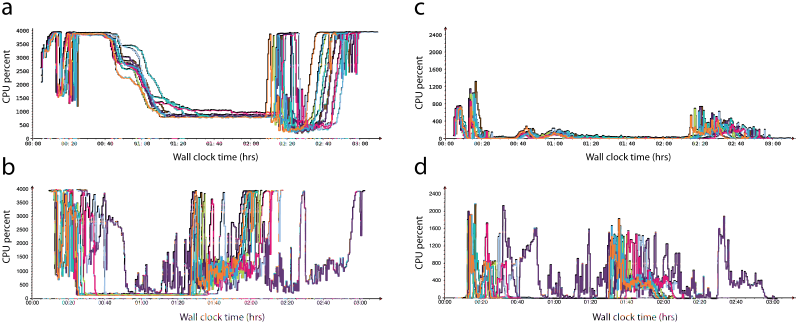
Comparison of undedicated cloud computing performances. The panel includes results of Amazon Web Services Elastic MapReduce (EMR) on Amazon EC2 instances (panels a & c) versus Google Compute Engine (GCE)(panels b & d) for human genome alignment and variant calling. In this 40 node cluster the total CPU percent for CPU idle (a and b) and waiting for disk input/output (c and d) is displayed. Note the greater consistency in performance of Crossbow, though generally longer wall clock times for complete analysis, on EMR compared to GCE.

## RESULTS

Wall-clock time for complete mapping and SNP calling differed by 52.9% (95%CI: 27.5-78.2) and 53.5% (95%CI: 34.4-72.6) for *E.coli* and human genome alignment and variant calling, respectively, with GCE being more efficient than EMR. Table 1 displays the key metrics for data analysis using both services. The proportion of central processing unit (CPU) usage by Crossbow differed between platforms when aligning and SNP calling each genome, with GCE having better utilisation as the genome size increased. There was considerably more free memory on GCE for the smaller *E.coli* dataset and on EMR for larger human genome runs. The CPU idle percentage, the percentage of time where the CPU was idle without waiting for disk input/output (I/O), was greater on EMR for the human genome while CPU waiting for I/O (WIO) was considerably lower on the same platform. The CPU idle and CPU WIO percentages were both significantly higher on EMR for the *E.coli* genome. The cost of running this Crossbow pipeline on EMR and GCE also differed significantly (p<0.001), with the costs on EMR being 257.3% (95%CI: 211.5-303.1) and 173.9% (95%CI: 134.6-213.1) more expensive than GCE for *E.coli* and human assemblies, respectively. For ∼36x coverage of a human genome, at a current sequencing cost of ∼US$1000, the median cost for computation on GCE was US$29.81 (range: US$28.86 to US$45.99), whilst on EMR with a fixed hourly rate it was US$69.60 (range: US$69.60 to US$92.80).

**Table 1.** **Comparison of performance metrics for genomic alignment and SNP calling.** All times are presented as hr:min:sec and remaining metrics are shown as mean ± standard deviation. * Calculated by paired *t*-test.

| Metric | <u>*E.coli* Genome</u> |  |  | <u>Human Genome</u> |  |  |
| --- | --- | --- | --- | --- | --- | --- |
|  | EMR (n=10) | GCE (n=10) | p-value* | EMR (n=10) | GCE (n=10) | p-value* |
| Wall clock time (mean) | 0:46:30 | 0:31:50 | <0.001 | 2:58:24 | 2:14:12 | <0.001 |
| Pre-processing short reads time (mean) | 0:14:37 | 0:12:46 | 0.109 | 0:07:29 | 0:06:23 | 0.116 |
| Alignment with Bowtie time (mean) | 0:07:04 | 0:05:03 | <0.001 | 1:51:06 | 1:15:07 | 0.003 |
| Calling SNPs with SOAPsnp time (mean) | 0:05:05 | 0:02:51 | <0.001 | 0:35:31 | 0:29:31 | 0.033 |
| Post-processing time (mean) | 0:04:51 | 0:00:57 | <0.001 | 0:01:23 | 0:01:03 | <0.001 |
| CPU user (mean %) | 17.44 ± 1.30 | 22.31 ± 3.14 | <0.001 | 43.80 ± 1.87 | 58.05 ± 6.20 | <0.001 |
| CPU idle (mean %) | 72.75 ± 1.23 | 65.76 ± 4.63 | <0.001 | 47.48 ± 2.30 | 22.17 ± 3.14 | <0.001 |
| CPU wio (mean %) | 3.88 ± 1.06 | 0.70 ± 0.16 | <0.001 | 1.86 ± 0.19 | 4.54 ± 1.82 | 0.001 |
| Bytes in (MB/sec) | 1.15 ± 0.09 | 2.12 ± 0.42 | <0.001 | 1.58 ± 0.07 | 2.00 ± 0.19 | <0.001 |
| Memory free (GB) | 2.19 ± 0.13 | 6.17 ± 0.42 | <0.001 | 0.91 ± 0.07 | 0.70 ± 0.03 | <0.001 |

Although runtime variability was inevitable and present in both platforms when assembling each genome, GCE had a considerably greater variability with the larger human genome compared to EMR (coefficient of variation (COV)_EMR_=4.48% vs COV_GCE_=16.72%). We identified a single outlier in run time on GCE during the human genome analysis. This occurred due to the virtual cluster having a slower average network connection (1.55MB/s compared to the average of the other GCE clusters of 2.02MB/s) and a higher CPU WIO percentage than the average for the other GCE runs (9.56% versus 3.52%). The variation in cluster performance likely reflects an increase in network congestion amongst GCE servers.

Runtime predictably is an important issue in undedicated cloud computing. The existing workload of the cloud at the time of service usage is one of the main determinants of variability in runtime of undedicated services.[12] In our benchmarking, EMR was more consistent, though slower, in overall wall-clock time compared to GCE. This may suggest that GCE is more susceptible to server congestion than EMR; though service usage data is difficult to obtain.

## DISCUSSION

Our findings confirm that cloud computing is an efficient and potentially cost-effective alternative for analysis of large genomic datasets. Cloud computing offers a dynamic, economical and versatile solution for large-scale computational analysis. There have been a number of recent advances in bioinformatic methods utilising cloud resources,[4,9,13] and our results suggest that a standard genomic alignment is generally faster in GCE compared to EMR. The time differences identified could be attributed to the hardware used by the Google and Amazon for their cloud services. Amazon offers a 2.0GHz Intel Xeon Sandy Bridge CPU, whilst Google uses a 2.6GHz Intel Xeon Sandy Bridge CPU. This clock speed variability is considered the main contributing factor to the difference between the two undedicated platforms. It must also be noted that the resource requirements of Ganglia may have had a small impact on completion times.[11]

There are a number of technical differences between GCE and EMR, which are important to consider when running standard bioinformatic pipelines. Running Crossbow on Amazon Web Services was simplified by an established support service, which provides an interface for establishing and running Hadoop clusters (Supporting Information 1). In contrast, there is currently no built-in support for GCE in Crossbow (Supporting Information 2). The current process to run a Crossbow job on GCE requires users to complete various steps such as installing and configuring the required software on each node in the cluster, transferring input data onto the Hadoop Distributed File System (HDFS), downloading results from the HDFS and terminating the cluster on completion. All of these steps are automatically performed by Crossbow on EMR. Python scripts offering similar functionality for GCE that Crossbow provides for EMR were created and are available (https://github.com/hewittlab/Crossbow-GCE-Hadoop).

While our findings confirm that cloud computing is an attractive alternative to the limitations imposed by the local environment, it is noteworthy that better performance metrics and lower cost were found with GCE compared to its established counterpart, Amazon’s EMR. Currently, a major limitation of these services remains at the initial transfer of large datasets onto the hosted cloud platform.[14] To circumvent this in the future, sequencing service providers are likely to directly deposit data to a designated cloud service provider, thereby eliminating the need for the user to double handle the data transfer.[15] Once this issue is resolved, it is foreseen that demand for these services is likely to increase considerably, given the low cost, broad flexibility and good customer support for cloud services.[15] The development of additional tools specific to genomic analysis in the cloud, which offer flexibility in choice of providers, is clearly required.

## METHODS

### Datasets and Analytical Pipeline

We benchmarked two platforms by a single job that completed read alignment and variant calling stages of next generation sequencing analysis simultaneously on two independent cloud platforms. To investigate the impact of data size on undedicated cluster performance, one small (*Escherichia coli* CC102 strain (3 GB SRA file; Accession: SRX003267) and one large (a Han Chinese male genome (142 GB Fastq files; Accession: ERA000005) publicly available genomic dataset was selected for analysis.[8,10] For each job in this experiment, a parallel workflow was designed using Crossbow. This workflow included the following four steps: (1) Download and conversion of files; (2) Short read alignment with Bowtie; (3) SNP call with SOAPsnp; and (4) Combination of the results (Supporting Information Figure 2). Crossbow was the preferred genetic annotation tool in this experiment, as it has built in support for running via Amazon’s EMR and Hadoop clusters.[16]

### Cluster construction and architecture

Instances were simultaneously established on Amazon’s EMR (http://aws.amazon.com/ec2/) and GCE (http://cloud.google.com/products/compute-engine.html). Undedicated clusters were optimized by selection of computational nodes as suggested for Crossbow.[9] Nodes with equivalent specifications were selected for each system (Supporting Information Table 1), these being c1.xlarge node in EMR and the closest specification node n1-highcpu-8 in GCE. For the *E.coli* genome, two nodes (one master and one slave) were used on each platform. On the other hand, for the human genome, the cluster was built with 40 nodes (one master and 39 slaves). As GCE did not provide any included storage for each instance, a 128GB drive (the default storage quota provided by GCE) was added for each node. This was at the additional cost of $0.04/GB/Month or $0.000056/GB/Hour (Jan to June 2014).

Each cluster was run using Apache Hadoop, an open-source implementation of the MapReduce algorithm.[17] MapReduce was used to organise distributed servers, manage the communication between servers and provide fault tolerance allowing tasks to be performed in parallel.[18]

To explore the effect of network activity differences between the platforms, each job was run simultaneously; same day (including weekdays and weekends) and same time. Detailed description of the set up and scripts to run the jobs can be found in Supporting Information Section 1 and 2.

### Cluster Monitoring

In both EMR and GCE, multiple components of cloud infrastructure including CPU utilisation, memory usage and network speeds were monitored and recorded for each node using Ganglia. The default setting of Ganglia for distributing incoming requests is multicast mode; however, since EMR and GCE environments do not currently support multicast Ganglia, it was configured in unicast mode (Supporting Information Figure 3). The metric output files constructed in .rrd format were converted into .csv format with a Perl script (Supporting Information Section 3). For comparison between performance and costs between platforms, the Student t-test was undertaken using the statistical software R (R Foundation for Statistical Computing version 3.0.2; http://www.r-project.org/). In the analysis, cost of each run was calculated using current pricing (June 10^th^ 2014); however, all *E.coli* runs and one human genome run were performed prior to a recent decrease in price on both platforms.

## ACKNOWLEDGMENTS

This work was supported by funding from the BrightFocus Foundation, the Ophthalmic Research Institute of Australia and a Ramaciotti Establishment Grant. CERA receives operational infrastructure support from the Victorian government.

## SUPPLEMENTARY MATERIAL

### Supplementary Section 1: Uploading data and setting up an Amazon Web Services Elastic MapReduce (EMR) cluster

Crossbow on EMR supports importing data referenced from the manifest file from HTTP, FTP, HDFS, and S3. We uploaded the input data (available from: ftp://public.genomics.org.cn/BGI/yanhuang/Rawdata2) into a S3 bucket in order to have the maximum potential download speed and ensure that the cluster was not sitting idle while the inputs were downloaded from a remote FTP server. The process used to upload the files from the remote FTP to our S3 bucket was as follows:

1. Create a micro instance on EC2, ensuring the instance has enough storage space for the whole dataset.
2. SSH into the instance using the given hostname,
3. (Optional) Install the screen package, and create a screen session. This allows you to close the SSH connection and allow the processes to continue. This is done via: sudo apt-get install screen Note: a screen session can be reconnected by: screen –r
4. Download the data from the remote FTP server: wget -r ftp://public.genomics.org.cn/BGI/yanhuang/Rawdata2
5. Upload the data to the S3 bucket s3cmd put --recursive /path/to/data/ s3://<YOUR-bucket>/path/to/save/data

The time taken to download the 142GB human genome data from the FTP server was 10:35:49 and the time taken to upload to the S3 bucket was 2:41:30, giving a total transfer time of 13:17:19.

The Ganglia install scripts provided by Amazon were using an out-of-date version of the software, which did not allow the metric data to be exported. As such these install scripts were updated to use the current version of Ganglia and Ganglia-web (3.6.0 and 3.5.10 respectively) and reinstalled. Ganglia was configured to use unicast mode as EMR prevented use of multicast.

Crossbow provided a faulty file in their S3 bucket that consistently caused the final step on EMR to fail. The file was hosted on our development bucket and Crossbow was modified so it was pointing to the S3 bucket. The link in the source code of Crossbow was altered and should now be corrected in the official Crossbow bucket.

### Supplementary Section 2: Scripts for configuration and running jobs on Google Compute Engine (GCE)

Crossbow provides an option that sets up and configures an EMR cluster with the required software. Although no such support is available for GCE, Crossbow supports use of a Hadoop Cluster that can be implemented in GCE. Google have released scripts for creating a Hadoop cluster on the GCE services (available from: https://github.com/GoogleCloudPlatform/solutions-google-compute-engine-cluster-for-hadoop). However, these scripts needed to be modified as we required some additional software to be installed on each node of the cluster. These modifications can be found at https://github.com/hewittlab/Crossbow-GCE-Hadoop. The modifications perform the following additional steps when setting up the cluster:

1. Create Ganglia configuration files and Hadoop configuration files specifically for the hostnames and IP addresses assigned to each of the nodes in the cluster.
2. Upload the configuration files to the Google Storage Bucket.
3. Install the required software on each of the nodes in the cluster, and place the configuration files from step 1 into their appropriate locations.

To run Crossbow on GCE, we initially uploaded the input reference Jar and the input manifest file to the HDFS. Given that these files were to be used for multiple runs and to avoid re-upload these files multiple times this was performed via the Google Storage Bucket. Once completed, the cluster was started. The input files were then downloaded from the bucket, onto the master node and then into the HDFS. Crossbow for Hadoop was then run with the appropriate parameters for the current cluster and input files, with the output files being copied from the HDFS to the bucket on completion. Each cluster was terminated upon completion of the workflow.

To reduce the repetition of steps required to set up the GCE cluster for each experimental run, an additional script, run_crossbow.py was created. While this script simplified the process to a level that was comparable to running Crossbow for EMR on the command line, it also raised several other issues. For example, access permission to the RRD directory on each node was made available to all user as required by Ganglia using ‘chown’ command.

Hadoop is designed to manage multiple failures and prevent a job from completely failing within a cluster. The default configuration of Hadoop stops the cluster immediately in case of a single failure. To prevent random task failures stopping the GCE cluster, we increased the threshold number of failed tasks allowable within the cluster.

When running Hadoop on GCE, a storage disk space had to be allocated to the instance at the time of its creation. Moreover, when creating a disk based off a Debian image, the available disk space was limited to 10GB regardless of the specified disk size. Linux partitions were edited to utilize the full size of the disk. A snapshot of the storage disk was created in order to avoid repartitioning of the disk during node creation. These modifications were updated further in the release version to reduce the setup time.

By default, Google only allows access to a limited number of resources per project. Since use of large cluster recommended in Crossbow runs, we applied for access to additional number of cores, IP addresses and total disk space in the region of our choice through an online form provided on Google Developers Console.

On January 9^th^ 2014, Google depreciated the v1beta15 application programming interface (API) and disk used for temporary storage was replaced with a persistent disk approach. This change caused problems in our initial cluster setup. Therefore, setup scripts were altered in order to generate a disk for each node prior its creation and to delete these disks upon termination. To point our code to the new API, the cluster creation scripts were modified based on a revised version of the Google’s code. However, later on we rewrote our changes into the updated Google scripts which are now available in our repository.

Finally, it is important to configure the number of Crossbow tasks appropriately to ensure complete node usage. The following arguments should be used with Crossbow on GCE: --cpus (number of cores per instance); and --instances (number of instances). It is important to note that these arguments are applicable to Hadoop in a non-EMR environment.

### Supplementary Section 3: Transformation of metric outputs from .RRD to .CSV format

The program rrd2csv.pl was used to convert RRDs to CSVs. It is available from https://code.google.com/p/rrd2csv/. Its usage is:

perl rrd2csv.pl -s “17/12/2013 12:00” -e “17/12/2013 13:00” file.rrd

To store the output as a csv, add “> file.csv” to the end (i.e perl rrd2csv.pl -s “17/12/2013 12:00” -e “17/12/2013 13:00” file.rrd > file.csv)

The -s and -e flags are representing the times that one adds the csv, where -s is the start time and -e is the end time. So in the example above, we are exporting from today at 12, to today at 1pm. These times should be when the job ran.

**Supplementary Figure 1.**
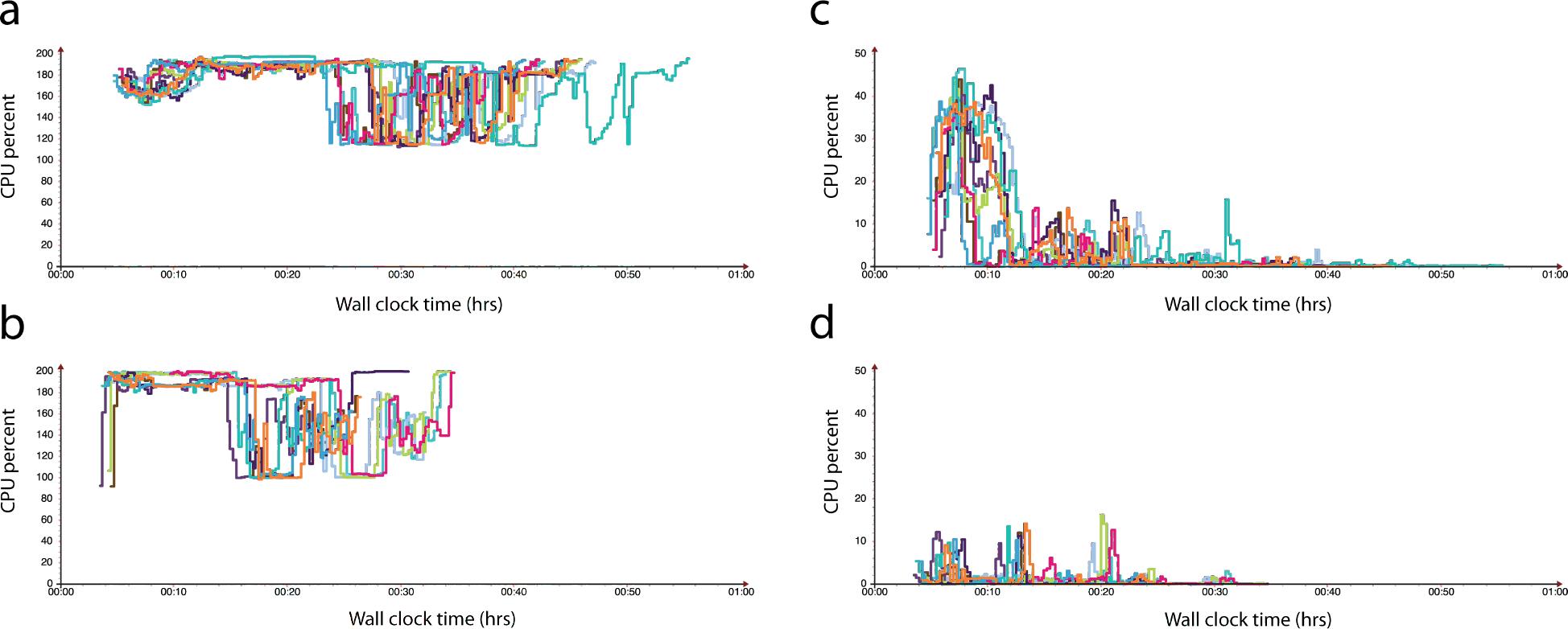
Comparison of undedicated cloud performance of Amazon Web Services Elastic MapReduce (EMR) on Amazon EC2 instances (panels a & c) versus Google Compute Engine (GCE)(panels b & d) for *E.coli* genome alignment and variant calling. In this two node cluster the total CPU percent for CPU idle (a and b) and waiting for disk input/output (c and d) is displayed. Note the shorter wall clock times for complete analysis on GCE compared to EMR.

**Supplementary Figure 2.**
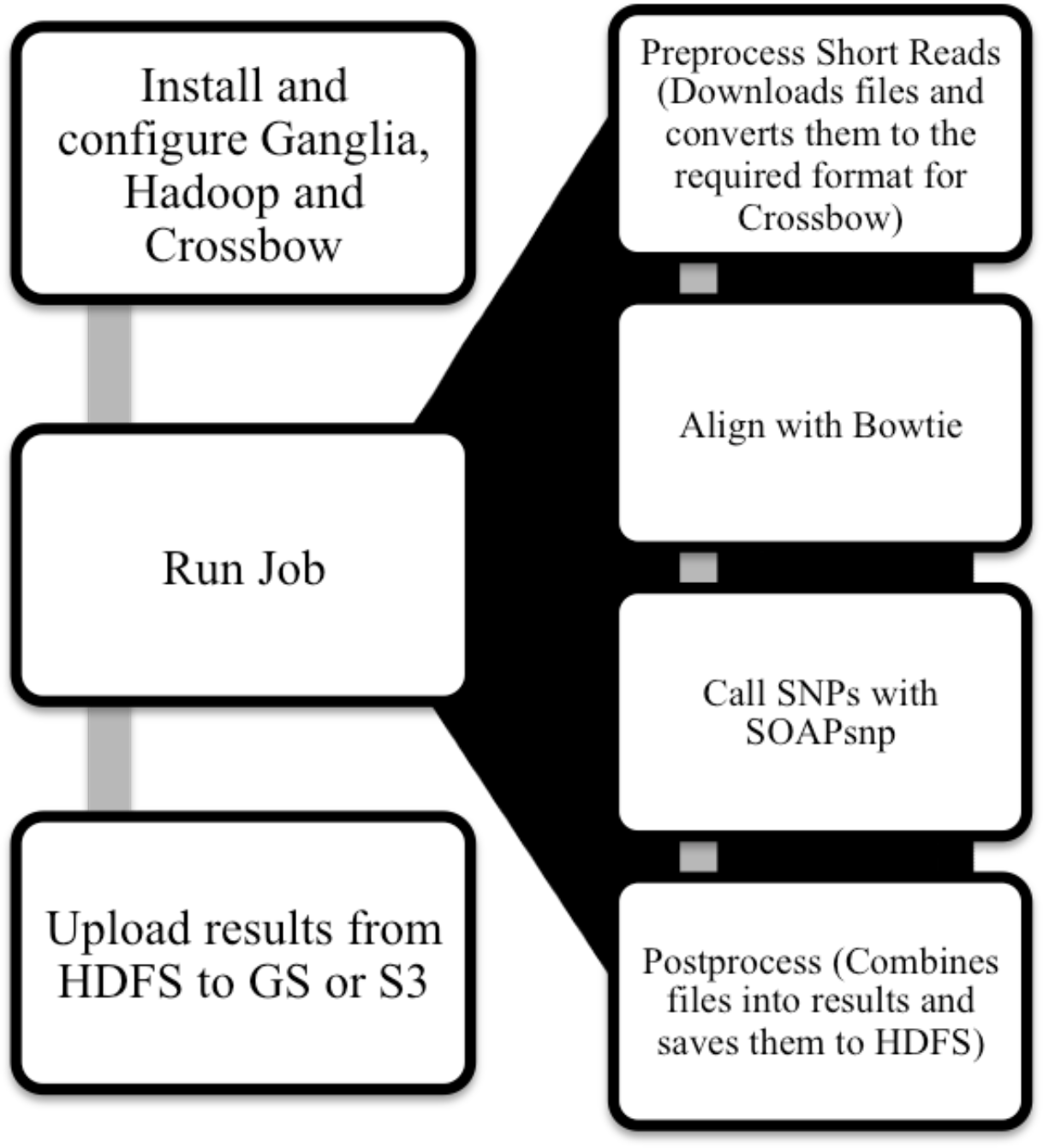
**Analytical pipeline demarcating each step required to complete the Crossbow job in the cloud.**

**Supplementary Figure 3:**
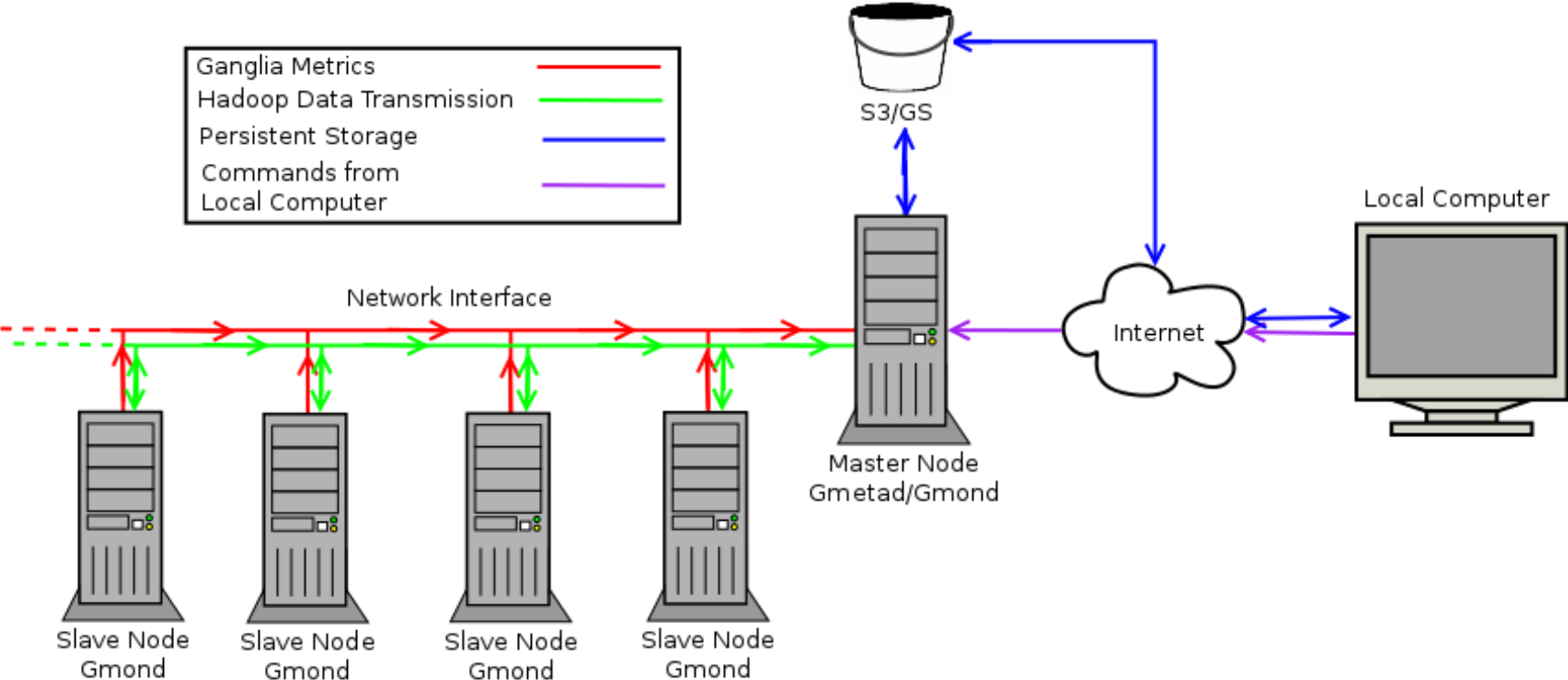
Directions and types of network transfers in our cloud-computing model. There are a variety of different network transfers between the nodes for each of the services in use in our model. Hadoop requires a bidirectional transmission of data between the master node and the slave nodes. This is required to coordinate the parallel processing of the cluster, and to allow for data transfer between nodes. Ganglia uses a unidirectional connection from the slave nodes to the master node to transfer the recorded metrics for storage and visualization. The persistent storage (provided by Amazon S3 (Simple Storage Service) or Google Storage, or an alternative method such as an FTP server) is accessed via the master node. The master node uses it to download input files for Crossbow, such as the manifest file and the reference Jar, and to use for persistent storage of the results of the Crossbow job as the instances destroy their storage on termination. Our local computer can also access the persistent storage via the Internet to allow access to upload the input files, or to download the results. The local computer needs to access the master node to initiate Crossbow. In EMR, this is replaced by a web interface and a JavaScript Object Notation Application Programming Interface (JSON API). In GCE, the user is required to remotely log in via Secure Shell (SSH) to commence the job.

**Supplementary Table 1:** Specification of used computational nodes for each system.

|  | **Virtual**<br>**Cores** | **Memory**<br>**(GB)** | **Included**<br>**Storage (GB)** | **Price**<br>**(USD/Hour)<sup>^</sup>** |
| --- | --- | --- | --- | --- |
| **Amazon Elastic Compute**<br>**Cloud (EC2) + Elastic**<br>**MapReduce (EMR)**<br>**[c1.xlarge]** | 8 | 7 | 4x420 | \$0.640 |
| **Google Compute Engine**<br>**[n1-highcpu-8]** | 8 | 7.2 | 0# | \$0.352 |

<sup>^</sup> Date accessed: April to June 2014; prior to this period, pricing was \$0.700 and \$0.520 in Amazon and Google respectively.

